# Motor unit behaviour adaptations across the lifespan: sex differences in young, middle-aged and old adults

**DOI:** 10.64898/2025.12.19.695460

**Authors:** Marco Carbonaro, Marta Boccardo, Clarissa M Brusco, Andrea M Pilotto, Rebecca Re, Fulvio Lauretani, Simone Porcelli, Martino V Franchi, Alberto Botter

**Author notes:** These authors share senior authorship (Martino V Franchi, Alberto Botter).

## Abstract

Ageing is associated with neuromuscular decline, and emerging evidence suggests that sex may influence the time course of motor unit adaptations. This study examined age- and sex-related differences in motor unit firing behaviour across young (YG), middle-aged (MA), and older adults (OLD), by integrating high-density EMG motor unit analysis with muscle morphology and daily physical activity measurements.

The analysis of single motor unit activity during submaximal isometric contractions of the vastus lateralis revealed that older adults had lower firing rates and a reduced capacity to modulate discharge frequency during force-increasing contractions. In the YG and MA groups, females showed higher motor unit firing rates and variability than males, while in OLD these sex differences were no longer present. Females also demonstrated a steeper decline in firing rate modulation between MA and OLD. Reductions in muscle cross-sectional area and thickness were similar between sexes. Physical activity levels declined with age in both sexes.

These findings reveal distinct, sex-specific trajectories of neuromuscular ageing, with females showing greater motor neuron function decline between MA and OLD, in the absence of sex-related differences in the rate of morphological deterioration. The attenuation of sex differences in older age suggests a convergence of neuromuscular profiles with ageing. While physical activity may contribute to the observed sex-specific patterns, other mechanisms related to hormonal shifts warrant further investigation. These insights underscore the importance of considering age and sex in the study of motor control and in the development of targeted interventions to preserve muscle function across the lifespan.

## INTRODUCTION

Ageing is a physiological process characterised by a progressive loss of muscle mass (Larsson et al., 2019), strength (Delmonico et al., 2009) and power (Tieland et al., 2018), accompanied by a decline in neuromuscular function (Hunter et al., 2016). Extensive evidence indicates that the nervous and musculoskeletal systems deteriorate in older adults, leading to several dysfunction in motor coordination and of the excitation–contraction coupling (Janssen et al., 2002). Ageing is linked to an adaptive remodelling of the neuromuscular system (Jones et al., 2022), which, however, still deteriorates from structural integrity and functional capacity perspectives (Drey et al., 2014; M. Piasecki et al., 2018)., with significant consequences for motor performance (Long & Pavalko, 2004; Wilkinson et al., 2018).

Relevant components of the neuromuscular system, such as motor neurons, neuromuscular junctions, and muscle fibres, undergo significant changes with advancing age, leading to modifications in motor unit behaviour across the lifespan (Guo et al., 2024; Roos et al., 1997; Sarto et al., 2024). One of the primary neurophysiological mechanisms associated with ageing is a progressive loss of motor neurons, often accompanied by instability at the neuromuscular junction (Drey et al., 2014; Erim et al., 1999). Between ∼50 and 60 years of age the body attempts to counteract the denervation of muscle fibres through reinnervation by surviving motor neurons, producing larger motor units and clustering of fibres (Lexell et al., 1988; Lexell & Downham, 1991). As ageing progresses these denervation-reinnervation cycles become increasingly inefficient (Brown, 1972; Brown et al., 1988; M. Piasecki et al., 2018). Beyond ∼75 years, denervation outpaces reinnervation, leading to a net reduction in muscle fibres, especially type II fibres (Deschenes, 2004; Horwath et al., 2025; Lexell et al., 1988). Neural changes therefore manifest not only as fewer motor units but also as reduced motor unit firing rates and increased synchronization, which could be seen as an effect of a reduced capacity of differentiating firing coding between motor units (Hepple & Rice, 2016; Nelson et al., 1984; Orssatto et al., 2022, 2023; Roos et al., 1997). In addition, early-recruited motor units increase their firing rate to a lesser extent during rising-force contractions in older individuals (Girts et al., 2020; Watanabe et al., 2016). Collectively, these modifications contribute to the decline in muscle force and endurance observed in older adults (Klass et al., 2008; Mani et al., 2018).

In this context, sex differences in motor unit characteristics, such as firing rate, recruitment threshold and discharge variability, have already been documented (Inglis & Gabriel, 2020, 2021; Jenz et al., 2023; Lulic-Kuryllo & Inglis, 2022; Marsala et al., 2025; Nishikawa et al., 2024; Olmos et al., 2023; Parra et al., 2020; Yacyshyn et al., 2025). Young women tend to discharge low-threshold motor units at higher rates during steady, moderate, isometric contractions, whereas men may exceed women firing rates at maximal intensities (Inglis & Gabriel, 2020, 2021; Marsala et al., 2025; Olmos et al., 2023). Women also show larger estimates of persistent inward currents, hinting at intrinsic motoneuronal differences (Jenz et al., 2023). Interestingly, in highly active masters athletes, while motor unit remodelling occurs similarly in both sexes only women exhibit progressive slowing of motor unit discharge with advancing age (J. Piasecki et al., 2021). Conversely, in less-active older cohorts, females maintain higher motor unit firing rates than age-matched males at a given relative torque yet still present smaller muscle cross-sectional area and poorer force steadiness (Guo et al., 2023). Taken together, these findings may suggest that differences in muscle phenotype and lifelong physical-activity patterns interact to produce sex-specific adaptations of motor unit behaviour across the adult lifespan.

In this scenario, much of our current understanding of motor unit firing properties in ageing has been derived from studies that either exclusively sampled men or pooled the sexes (Lulic-Kuryllo & Inglis, 2022). Given the growing evidence that males and females display distinct neuromuscular ageing trajectories—whether in terms of muscle-mass loss, firing-rate slowing or hormone-related modulation (Guo et al., 2023; Inglis & Cabral, 2025; O’Bryan et al., 2025)—it has become more and more important to differentiate the age stratification by sex. Accordingly, the present study aims to characterize sex differences in motor unit recruitment and firing properties at two critical stages of later life: middle-aged and older adulthood. These cohorts are compared against young adults to determine how motor unit behaviour diverges before overt neuromuscular decline sets in. Clarifying these variations is essential for designing interventions that preserve muscle function and physical performance in both women and men as the population ages (Sundberg, 2025). Expanding motor unit research to include balanced male–female samples is therefore not only scientifically rigorous but also central to developing effective, sex-sensitive health strategies (Graf et al., 2023).

## METHODS

This study is part of a longitudinal project named TRAJECTOR-AGE (Lauretani et al., 2025) aiming to investigate the 2-year time course of neuromuscular, structural, and metabolic changes and their interplay occurring with ageing. All the procedures used in this study comply with the Declaration of Helsinki and are registered at http://clinicaltrials.gov (NCT06168591) and were approved by the AVEN Ethical Committee (Emilia Romagna region, Italy) on July 5, 2022 (protocol #28022; study ID 283/2022/SPER/UNIPR).

### Subjects

Ten young (YG) adults, forty-six middle-aged (MA) adults, and thirty-five old (OLD) adults were enrolled in this study. All participants characteristics are reported in Table 1. The young group was composed of healthy participants from 20 to 30 years old with no neurological and musculoskeletal impairments or disease. The MA and OLD community-dwelling, mobile, and cognitively unimpaired participants were enrolled based on the following criteria: Mini-Mental State Examination score ≥ 24, gait speed > 0.8 m/s (assessed over 4 meters), Short Physical Performance Battery score > 9, Timed Up-and-Go test < 20 seconds, and grip strength > 27 kg for males or > 16 kg for females. Exclusion criteria comprise neurological disorders (including stroke), diabetes, late-stage cancer, severe chronic kidney disease, severe liver insufficiency, severe cardiac disorders, major recent heart surgery, and severe osteoarthritis. All participants signed written informed consent for the study after a detailed explanation of the purposes and procedures.

**Table 1.** Anthropometric measurements of the 91 participants divided by sex and age.

|  | Young |  | Middle-Age |  | Old |  |
| --- | --- | --- | --- | --- | --- | --- |
|  | Male (n = 5) | Female (n = 5) | Male (n = 17) | Female (n = 29) | Male (n = 25) | Female (n = 11) |
| **Anthropometry** |  |  |  |  |  |  |
| Age (years) | 26.4 ± 2.5 | 25.4 ± 2.2 | 58.7 ± 3.3 | 57.8 ± 3.2 | 77.3 ± 4.7 | 77.1 ± 4.1 |
| Height (cm) | 176.0 ± 6.7 | 167.4 ± 10.4 | 178.2 ± 5.4 | 163.7 ± 5.4 | 171.4 ± 8.1 | 155.6 ± 6.2 |
| Body mass (kg) | 67.4 ± 6.2 | 58.0 ± 13.6 | 86.4 ± 12.3 | 64.4 ± 10.6 | 80.3 ± 12.1 | 62.9 ± 9.4 |
| BMI (kg/m<sup>2</sup>) | 21.8 ± 2.2 | 20.5 ± 3.0 | 27.2 ± 3.8 | 24.0 ± 3.7 | 27.2 ± 2.6 | 26.1 ± 4.7 |

### Experimental protocol

High-density electromyographic (HDEMG) signals and force profiles were detected from the vastus lateralis (VL) during varying-force, isometric contractions (Figure 1). At the beginning of the experimental procedure, the muscle architecture of all participants was assessed to document the anatomical and morphological characteristics of the analysed muscle. Ultrasound images of VL muscle were acquired with participants lying relaxed supine, maintaining a hip angle of 160° (see *Ultrasound images recordings* for detailed procedures).

**Figure 1.**
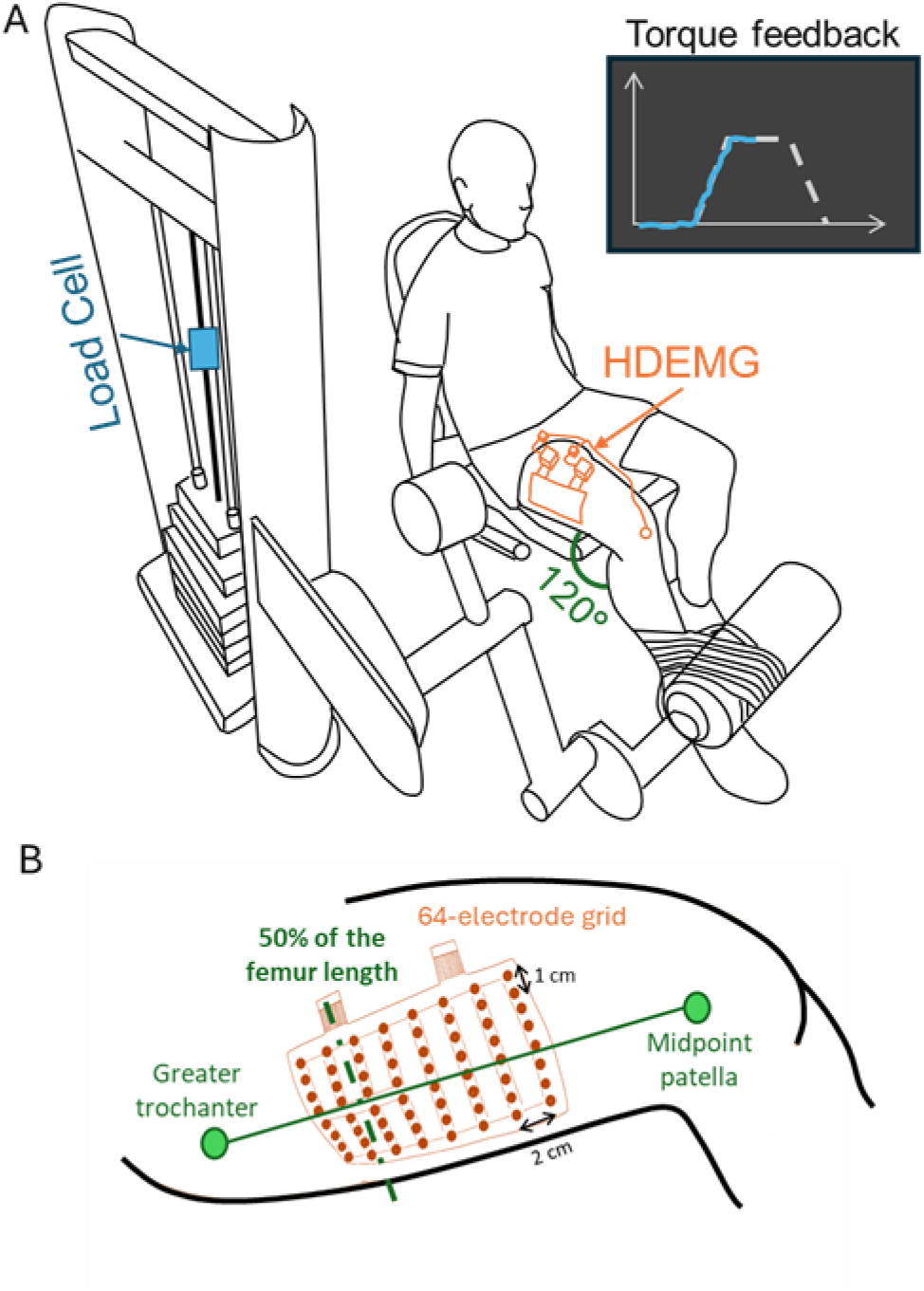
Experimental Setup. A) Instrumented leg extension machine for isometric contractions at a knee angle of 120° (where 180° is the full extension). A monitor displayed the time course of the recorded torque and the target torque level for visual feedback. B) Position of the 64-electrode HDEMG grid over the VL muscle at 50% of the femur length measured as the distance between the greater trochanter and the distal boundary of the lateral femoral condyle.

Following ultrasound imageing recordings the neuromuscular evaluation was conducted. Participants were seated on a leg extension machine (Figure 1A, BH L010 Leg Extension, BH Fitness-EXERCYCLE, S.L., Vitoria-Gasteiz, Spain) with the right leg secured to the pushing arm at a knee angle of 120° (where 180° is the full extension). The knee extension machine was instrumented to measure force through a load cell under isometric conditions (see *Force* recordings for details). The time course of the recorded torque and the target torque level were displayed to the participants on a monitor as visual feedback. During the warm-up phase, participants were instructed on the use of the visual feedback by performing contractions of varying intensities, ranging from low to high. Subsequently, each participant performed three maximal voluntary contractions (MVCs), each sustained for 5 seconds. The highest recorded torque value from these trials was designated as the MVC torque and served as a reference for subsequent contractions. Isometric knee extensions of the VL muscle were then performed while force and HDEMG signals were recorded (see *HDEMG recordings* for detailed methodology). Participants were required to match a trapezoidal force profile displayed on the feedback screen. The trapezoidal profile consisted of three phases: (i) a ramp phase from 0% to 30% or 50% of MVC with a slow force increase (2% MVC/s); (ii) a plateau phase lasting 15 seconds at either 30% or 50% of MVC; and (iii) a descent phase from 30% or 50% MVC back to 0% at the same rate of force change as the initial ramp phase. Participants performed one trapezoidal contraction at 30% MVC during the training phase. Following this, four contractions were recorded: two trapezoidal contractions with a plateau at 30% MVC, two with a plateau at 50% MVC. All contractions were separated by two minutes of rest.

At the end of the experimental protocol, the participants were also monitored during one week with an actigraphy to measure the level of general physical activity during normal life (see *Physical activity* assessment paragraph for detailed methodology).

#### Ultrasound images recordings

Ultrasound imageing (Esaote, Genoa, Italy) was performed by two expert operators (CMB, MVF) using a linear probe (LA523, frequency range 6-12 MHz) to measure VL cross-sectional area (CSA), muscle thickness, and subcutaneous adipose tissue thickness. The assessments were performed with the probe placed at 50% between the greater trochanter and the distal boundary of the lateral femoral condyle (Franchi et al., 2019; Narici et al., 2021) using water-based conductive gel applied to the surface of the ultrasound probe, facilitating better acoustic coupling while minimizing contact pressure on the skin. CSA was assessed with the probe transversally aligned to the muscle using the extended-field-of-view. After identification of the border where the VL meets the rectus femoris, the probe was aligned and moved slowly and continuously in the medio-lateral direction to construct the image of the whole muscle (Monti et al., 2020). Images were also acquired with the probe placed longitudinal to the muscle to standardize the point in which muscle thickness and subcutaneous adipose thickness tissue were assessed. A snapshot was performed at 50% of the femur length and at 50% of the distance between the mid-and lateral ends of the VL. For the young individuals, only the snapshot of the VL (ArtUs, Telemed) was acquired with a 38.4 mm linear transducer (L12-5N40-A4 - frequency range 5-12 MHz).

#### Force recordings

The force exerted during knee extension contractions was measured with a load cell (MODEL TF 031, full scale 100 kg, sensitivity 2 mV/V, CCT Transducers, Torino, Italy) placed in series with the cable of the leg extension machine. The load cell output was amplified with a general-purpose acquisition system (GAM, ReC Bioengineering and LISiN, Politecnico di Torino, Turin, Italy) synchronized with the HDEMG amplifier. To provide real-time feedback on the force generated by the participant, a custom script developed in Matlab (R2022b, The MathWorks Inc., MA, USA) was used to display the force levels on a monitor during the experiment.

#### HDEMG recordings

Muscle activity of the VL was recorded using a 64-electrode grid (arranged in 8 columns with a 10 mm inter-electrode distance and 8 rows with a 20 mm inter-electrode distance), covering an area of 70 × 140 mm (Figure 1B) specifically designed for the study. Ultrasound imageing was employed to identify the direction of the muscle fascicles at 50% of the femur length. After skin preparation, the electrode grid was securely positioned over the belly of the VL muscle (Figure 1B), ensuring that 50% of the femur length was aligned between the second and third columns (from proximal to distal), with the rows aligned to the fascicles’ direction previously identified.

Monopolar HDEMG signals were detected and conditioned (bandwidth 10–500 Hz, gain 192), then sampled at 2048 Hz with 16-bit resolution using a wireless high-density electromyography acquisition system (MEACS, ReC Bioengineering and LISiN, Politecnico di Torino, Turin, Italy) (Cerone et al., 2019). Electrophysiological signals and torque data were synchronized during post-processing using common external pulses that were simultaneously recorded by both the HDEMG system and the torque measurement system (Cerone et al., 2022).

#### Physical activity assessment

Physical activity was assessed using the triaxial wrist-worn Actigraph GT9X Link (Actigraph, Pensalcola, FL, USA) accelerometer on the non-dominant wrist with a sampling frequency of 60 Hz. Participants were fitted with the accelerometer and instructed to wear it for the next 7 days, 24 h/day, in the free-living environment. After completing the 7-day data collection, participants returned the accelerometer to the clinical research centre.

### Data Analysis

#### Measurements of the VL muscle morphology

The captured images were subsequently analysed using ImageJ software (National Institutes of Health, USA). CSA was analysed using the freehand tool and determined as the maximum muscle area excluding aponeurosis; muscle thickness was determined as the vertical distance between superficial and deep aponeurosis; and subcutaneous adipose tissue thickness was determined as the vertical distance between the skin and the superficial aponeurosis.

#### Torque signal

Force signals were converted into torque by multiplying by the constant lever arm of the leg extension. Signals were then low-pass filtered at 100 Hz (4th-order Butterworth), for computing the torque steadiness between the start and ending points of the plateau phase (identified by further low-pass filtering the signal at 2 Hz). During this phase, torque steadiness was determined computing the coefficient of variation (CoV) as the percentage ratio between the standard deviation and the mean value (Enoka & Farina, 2021; Galganski et al., 1993).

#### HDEMG signals decomposition and editing

The HDEMG signals were bandpass filtered within the 20–400 Hz range using a 4^th^ order Butterworth filter. Following this preprocessing, the signals were decomposed into individual motor unit firing patterns using a validated method based on convolution kernel compensation (CKC) (Holobar et al., 2010; Holobar & Zazula, 2007). This decomposition was performed with a Matlab-based software (DEMUSE, System Software Laboratory, Faculty of Electrical Engineering and Computer Science, University of Maribor, Slovenia). The decomposition results were visually inspected by two expert operators (MC and MB), who conducted an editing of the firing patterns, following best practices recommended in the literature (Del Vecchio et al., 2020; Merletti & Muceli, 2019).

#### Analysis of motor unit firings behaviour

Motor unit behaviour was assessed by computing the following variables of interest: (i) the mean and the coefficient of variation (CoV) of the firing rate of all the decomposed motor units during the constant-torque phase at 30% and 50% of the MVC, (ii) the difference between the firing properties of motor units grouped for torque levels (from lower to higher threshold motor units), (iii) the firing rate modulation as a function of the exerted torque.

During the ramp-up phases of the torque, HDEMG signals were segmented into intervals corresponding to 10% of the MVC to identify groups of motor units recruited at different torque levels (Figure 2A). This segmentation resulted in three motor unit groups for the 30% MVC trials (Figure 2A) and five motor unit groups for the 50% MVC trials. For each motor unit group, the mean firing rates were computed for the torque interval in which they were recruited and for the following ones (Watanabe et al., 2016). Figure 2B shows an example for the 30% MVC contraction: the mean firing rate for first group (i.e. motor unit recruited within the 0-10% MVC interval) was evaluated across all three contraction intervals (0-10%, 10%-20%, and 20%-30% MVC). For the group of motor units recruited within the second interval (i.e. 10-20% MVC), the mean firing rate was computed only for the 10-20% and 20-30% MVC intervals. Similarly, this approach was applied to the 50% MVC contractions considering five motor unit groups and five contraction intervals (Watanabe et al., 2016).

**Figure 2.**
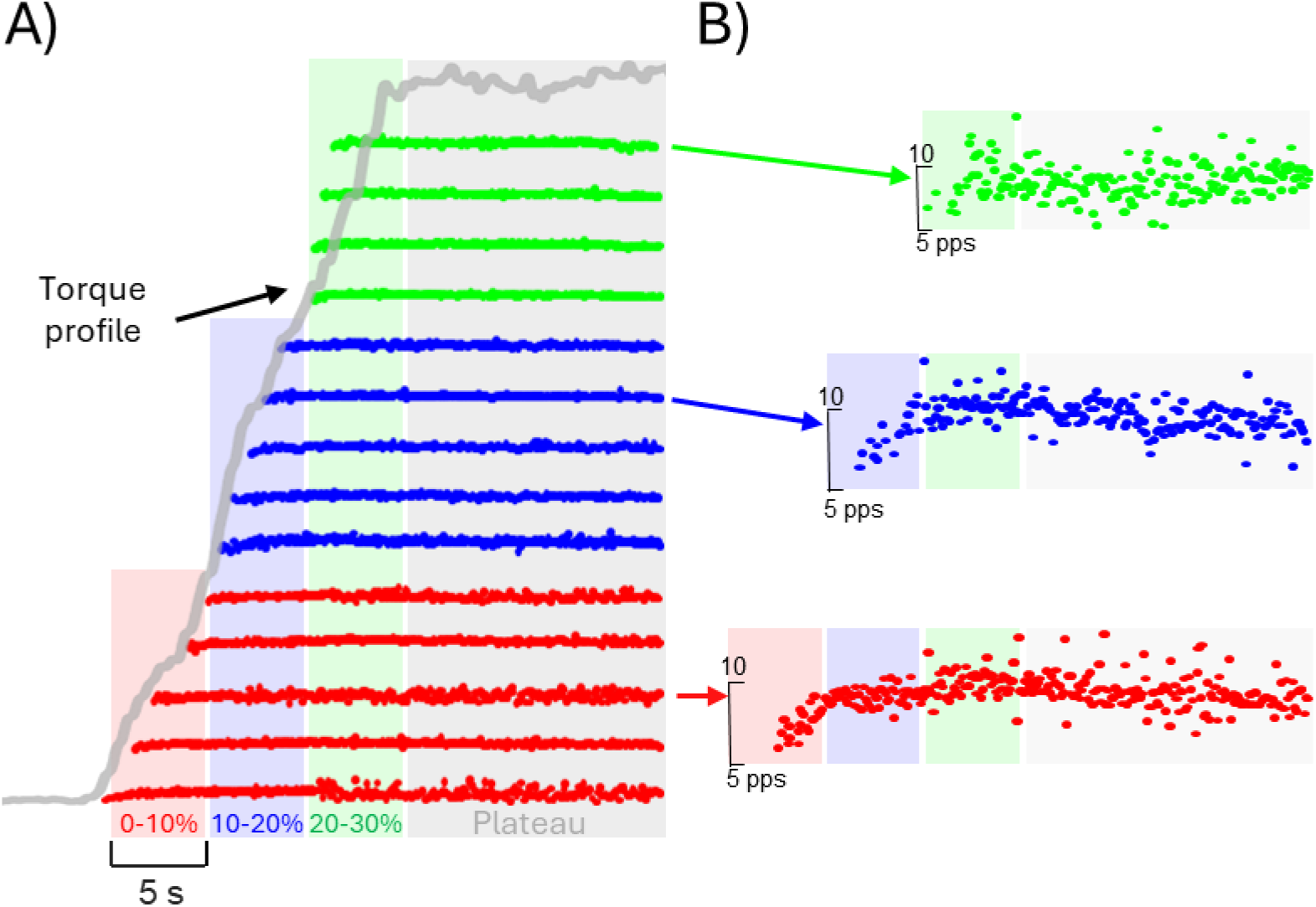
Example of decomposition results and analysis for a representative 30% MVC contraction. A) The ramp was segmented into intervals corresponding to 10% of the MVC to identify groups of motor units recruited at different contraction levels (red 0-10% MVC, blue 10-20% MVC, green 20-30% MVC).

Considering that the firing rate of a motor unit increases with the force level, this modulation was quantified for each motor units as the difference between the firing rates at the recruitment interval and the maximum torque (Δ firing rate). This analysis was performed for the 50% ramp and all the motor units were split in two groups, the early recruited (between 0 and 20% MVC) and the late recruited (between 20 and 40% MVC).

#### Measurements of physical activity

Accelerometer data were downloaded and pre-processed using the ActiLife software (version 6.13.4) to derive 1-min epoch activity counts. Only participants with ≥4 valid days of accelerometer data were included in the analysis. A valid day was defined as ≤10% missing data; non-valid days were excluded. We calculate total activity counts (TAC) per day and time spent in active states. TAC was derived from the sum of activity counts for each minute during a valid day. Each minute was labelled as active if the activity counts for that minute were ≥1853 or sedentary if they were <1853 (Koster et al., 2016). Data were processed with *R arctools package* (Karas et al., 2021).

### Statistics

The MVC torque, subcutaneous adipose tissue thickness, CSA, and muscle thickness of the VL muscle, as well as physical activity metrics (total activity and active time) and torque steadiness (CoV of torque during the plateau), were compared across age groups and sexes using two-way ANOVA. To account for body composition, MVC torque, CSA, muscle and subcutaneous thickness were further analysed using ANCOVA, with BMI included as a covariate. These analyses were conducted in R (version 4.3.1, R Foundation for Statistical Computing, Vienna, Austria).

The average firing rate (mean firing rate during the plateau), firing rate variability (CoV of firing rate during the plateau), and motor unit modulation (Δ firing rate) were compared across subject groups and sexes using either the Wilcoxon signed-rank test or the Mann–Whitney U test, depending on the outcome of the Shapiro–Wilk normality test applied to each variable distribution. The significance level (p < 0.05) was adjusted for multiple comparisons (Bonferroni). These analyses were conducted in MATLAB (R2022b, The MathWorks Inc., MA, USA).

Changes in firing rate of the same motor unit across successive contraction intervals within a single contraction (Force interval) were assessed using a mixed linear model, with Age, Sex, motor unit group, and Force interval as fixed effects, and Subject as a random effect. This analysis was conducted in R.

## RESULTS

### Muscle morphology

The analysis of muscle morphology across age and sex groups revealed significant differences, as reported in Table 2. There were significant differences in muscle CSA and muscle thickness between males and females, as well as between different age groups. Muscle thickness was significantly greater in males than females in all age groups (p < 0.05). Similarly, CSA was significantly larger in MA males compared to MA females (p < 0.001). When comparing for the same sex, MA exhibited a significantly larger CSA (p < 0.01) than the OLD. No significant differences in muscle thickness were observed between YG and MA groups for females and males (p > 0.05), while YG showed significant larger muscle thickness than OLD (p < 0.05) for both sexes. Subcutaneous adipose tissue thickness of females was larger than males for all age groups (p < 0.05) with no differences between ages (p > 0.05).

**Table 2.**
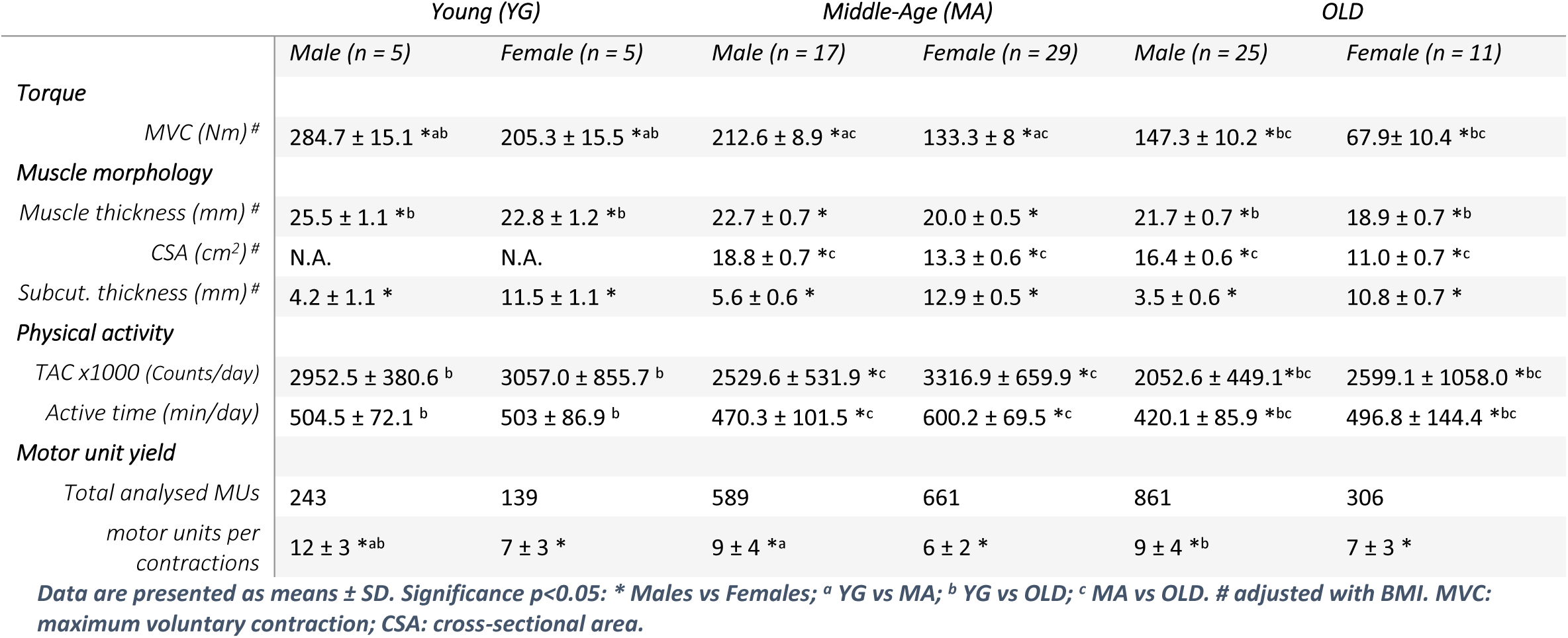
Results of the measurements of torque, muscle morphology, physical activity, and motor units.

|  | Young (YG) |  | Middle-Age (MA) |  | OLD |  |
| --- | --- | --- | --- | --- | --- | --- |
|  | Male (n = 5) | Female (n = 5) | Male (n = 17) | Female (n = 29) | Male (n = 25) | Female (n = 11) |
| **Torque** |  |  |  |  |  |  |
| MVC (Nm) # | 284.7 ± 15.1 *ab | 205.3 ± 15.5 *ab | 212.6 ± 8.9 *ac | 133.3 ± 8 *ac | 147.3 ± 10.2 *bc | 67.9 ± 10.4 *bc |
| **Muscle morphology** |  |  |  |  |  |  |
| Muscle thickness (mm) # | 25.5 ± 1.1 *b | 22.8 ± 1.2 *b | 22.7 ± 0.7 * | 20.0 ± 0.5 * | 21.7 ± 0.7 *b | 18.9 ± 0.7 *b |
| CSA (cm<sup>2</sup>) # | N.A. | N.A. | 18.8 ± 0.7 *c | 13.3 ± 0.6 *c | 16.4 ± 0.6 *c | 11.0 ± 0.7 *c |
| Subcut. thickness (mm) # | 4.2 ± 1.1 * | 11.5 ± 1.1 * | 5.6 ± 0.6 * | 12.9 ± 0.5 * | 3.5 ± 0.6 * | 10.8 ± 0.7 * |
| **Physical activity** |  |  |  |  |  |  |
| TAC x1000 (Counts/day) | 2952.5 ± 380.6 <sup>b</sup> | 3057.0 ± 855.7 <sup>b</sup> | 2529.6 ± 531.9 *c | 3316.9 ± 659.9 *c | 2052.6 ± 449.1 *bc | 2599.1 ± 1058.0 *bc |
| Active time (min/day) | 504.5 ± 72.1 <sup>b</sup> | 503 ± 86.9 <sup>b</sup> | 470.3 ± 101.5 *c | 600.2 ± 69.5 *c | 420.1 ± 85.9 *bc | 496.8 ± 144.4 *bc |
| **Motor unit yield** |  |  |  |  |  |  |
| Total analysed MUs | 243 | 139 | 589 | 661 | 861 | 306 |
| motor units per contractions | 12 ± 3 *ab | 7 ± 3 * | 9 ± 4 *a | 6 ± 2 * | 9 ± 4 *b | 7 ± 3 * |

Data are presented as means ± SD. Significance $p < 0.05$ : \* Males vs Females; <sup>a</sup> YG vs MA; <sup>b</sup> YG vs OLD; <sup>c</sup> MA vs OLD. # adjusted with BMI. MVC: maximum voluntary contraction; CSA: cross-sectional area.

### Physical activity

As reported in Table 2, we found significant effects for sex (p < 0.05) and age (p < 0.01) in both physical activity variables. MA and OLD comparison between sexes revealed higher TAC in females compared to males (p < 0.001) and, also the time spent active was higher in females (p < 0.001). No sex differences were found for both TAC and active time in YG (p > 0.06). Concerning age differences, the TAC and active time measured in OLD was significantly lower compared to MA and YG (both sexes p < 0.05). While YG males showed significantly higher TAC compared to OLD (p < 0.05).

### Torque measures

Males showed higher MVC torque compared to females in all age groups, with a significant reduction with age for both sexes (p < 0.05, Table 2). The CoV of the torque (torque steadiness) at 30% and 50% MVC contractions is shown in Figure 3A. At 30% MVC, females showed higher CoV of the torque (i.e. lower steadiness) than males in both MA and OLD groups (p < 0.05). Age-related differences were significant in females, with YG showing lower CoV than OLD (p < 0.05). At 50% MVC, no sex differences were observed, but CoV of the torque was significantly lower in YG females compared to MA and OLD (p < 0.05). Males showed a similar variability of CoV of the torque with age (at both 30% and 50% MVC, p > 0.3).

**Figure 3.**
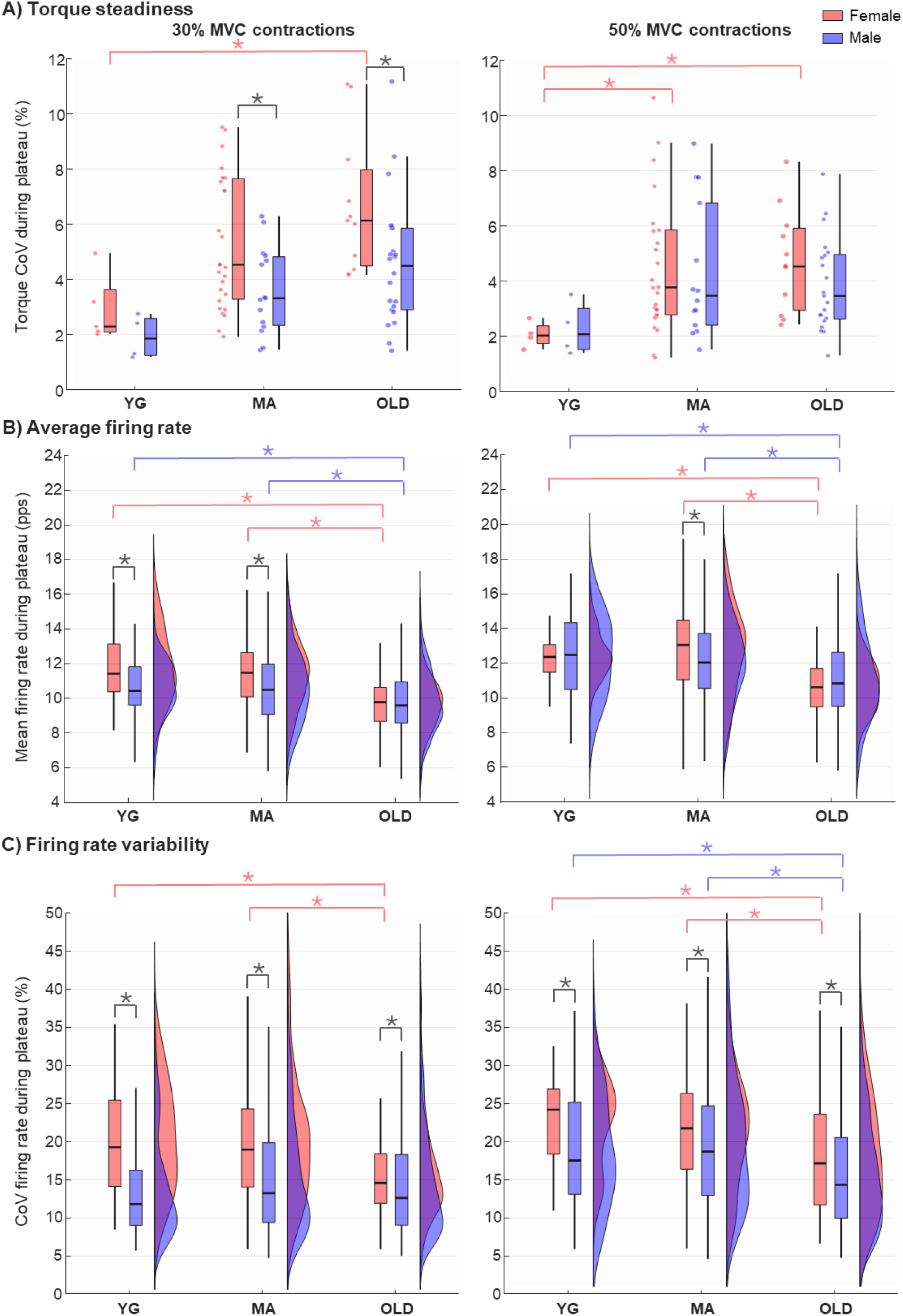
Results of torque steadiness, average firing rate, and firing rate variability. A) Boxplots and jitter plots of the coefficient of variation of the torque during the plateau phase of both 30 and 50% MVC contractions for all the subjects divided by age group and sex. B) Boxplots and probability density functions of mean firing rate, and C) coefficient of variation of the firing rate for all the motor units divided by age group and sex. Statistical significances (*p<0.05) are reported in black for comparison between sex and with the respective colour for comparison among age groups.

### Motor unit firing properties

The number of analysed motor units per contraction (Table 2) highlighted that higher motor units can be identified in males than females for all ages (p < 0.05) with the YG males showing the highest number of identification. Figure 3B shows the average firing rate of all the decomposed motor units during the plateau phase of the contraction at 30% (left panel) and 50% (right panel) MVC. Females showed higher firing rates than males in YG and MA (p < 0.001) but not in OLD (p = 0.95) at 30% MVC, while at 50% MVC this sex difference was observed only in MA (p < 0.01). For both sexes and contraction levels, the OLD group showed lower firing rates than YG (p < 0.001) and MA (p < 0.001). Figure 3C shows the CoV of the firing rate during the plateau for both 30% and 50% MVC. Females showed higher firing rate variability than males for all the age groups at both contraction levels (p < 0.001). OLD showed lower CoV of the firing rates than YG and MA for both sexes at 50% MVC (p < 0.001) and only for females at 30% MVC (p < 0.001). Figure 4A shows the modulation of firing rates during the ramp of the five groups of motor unit recruited at progressively increasing torques (10% MVC intervals) for the 50% MVC contractions. The firing rates of motor units increased with the force level in all groups of participants, but this modulation was less prominent in the OLD than in the YG subjects, particularly for the females. Statistical comparisons revealed that, at each contraction interval, earlier recruited motor units (e.g., group 1) fire at significantly higher rates than later recruited ones (e.g., groups 3–5) (p < 0.05 for multiple pairwise comparisons; see labels a–i). This difference is observed in YG and MA individuals, especially in males, where multiple significant comparisons across motor unit groups were evident at each force level. In contrast, in the OLD group (particularly females) the modulation becomes less prominent, and therefore fewer significant differences can be observed within each contraction interval. Figure 4B highlights this phenomenon by means of the Δ firing rate. For the early recruited motor unit group, significant sex differences were observed in MA and OLD (p < 0.01) but not YG (p = 0.7). MA females showed higher modulation than males (p < 0.01), while a reduced modulation of females than males (p = 0.3) was observed in OLD. No sex differences were observed for the late recruited motor units (p > 0.05). A trend in the reduction of modulation between ages was evident and was significant for both males and females and for all recruitment threshold intervals (p < 0.001), with the motor units in the OLD group showing a reduced modulation of the firing rate from the beginning to the end of the ramp.

**Figure 4.**
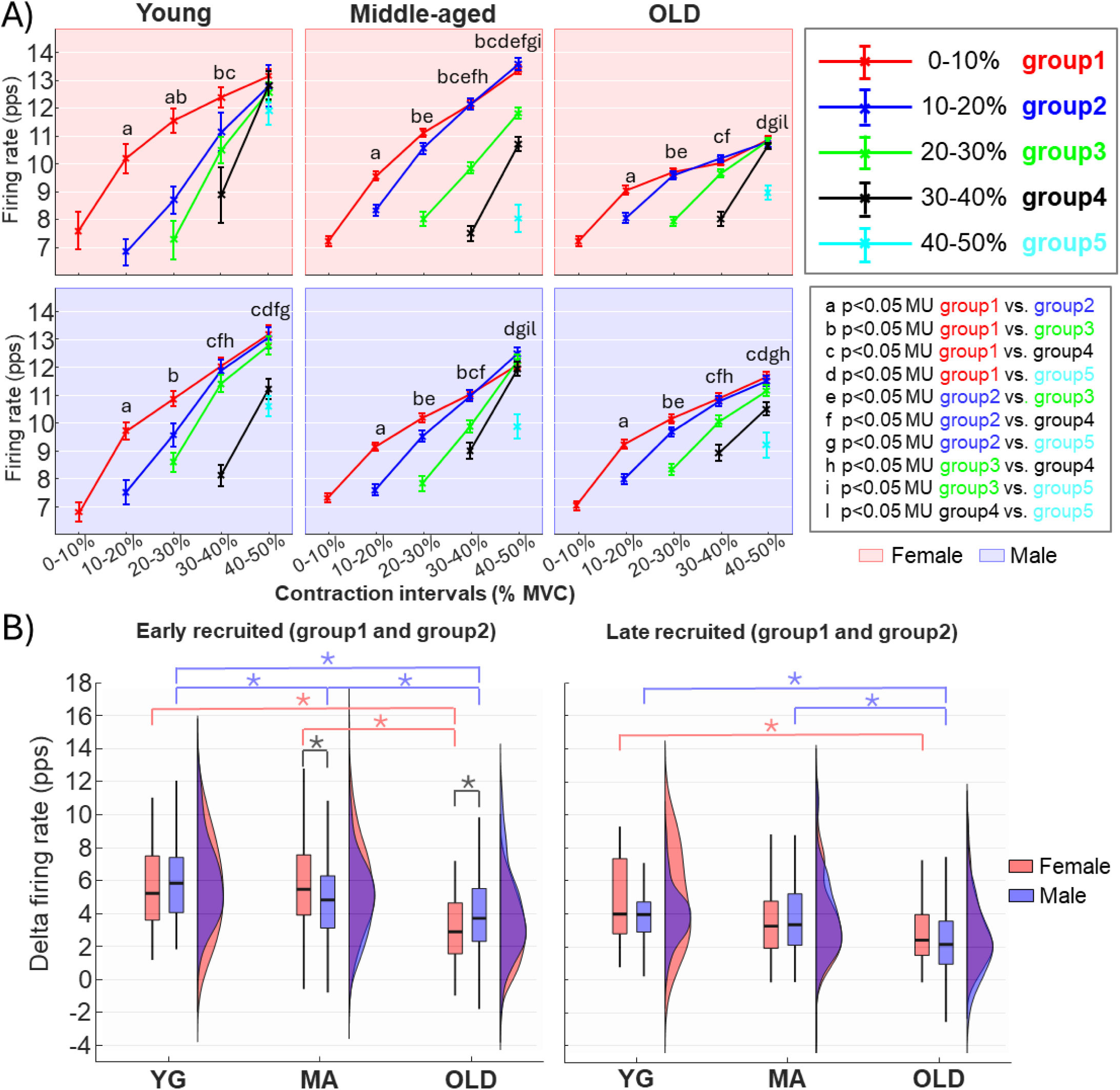
Firing rate modulation with increasing force. A) Modulation of firing rates during the ramp of the five motor unit groups recruited at different thresholds for the 50% MVC contractions. The upper panels belong to female divided among age groups, while the lower panels refer to male. All the motor unit groups were compared at all the contractions intervals using mixed linear models (see legend with significances for all the comparisons). B) Boxplots and probability density functions of the delta firing rate (difference between the average firing rates at the recruitment interval and at the maximum force) of motor units divided in two major group (early and late recruited) among different age groups and sexes. The early recruited group (left panel) consisted of the motor units recruited up to 20% MVC (group 1 and 2) while the late recruited group (right panel) consisted of the motor units recruited in the range 20-40% MVC (group 2 and 4). Statistical significances (*p<0.05) are reported in black for comparison between sex and with the respective colour for comparison among age groups.

## DISCUSSION

This study was designed to investigate motor unit behaviour across different stages of adulthood, with a particular focus on sex-related differences in neuromuscular ageing. The rationale stems from growing evidence that ageing affects neuromuscular system differently in males and females, yet few studies have examined age-related changes in parallel across multiple age groups. By integrating HDEMG-based motor unit analysis with the assessment of muscle morphology and daily physical activity, we aimed to provide an integrated view of the factors contributing to neuromuscular decline.

The main findings revealed a progressive change in motor unit firing properties with age, particularly a reduced discharge rate and an impaired ability to modulate discharge rate during increasing torque contractions. This decline was more pronounced in females transitioning from middle age to old age, suggesting a steeper deterioration in neural control mechanisms. In contrast, males exhibited a more linear decline from young to old age. Interestingly, while neural properties converged between sexes in older age, muscle morphology maintained consistent sex differences across all age groups. These findings, together with the evidence that physical activity is a critical modulating factor, suggest distinct morphological and neural ageing trajectories in males and females in the adult lifespan.

The first evidence in that the OLD group showed lower firing rates than YG and MA groups at both contraction levels, consistent with evidence of age-related declines in motor unit discharge rates (Orssatto et al., 2022). Moreover, the progressive change in motor unit firing properties with age was sex-specific. Females showed a more pronounced transition from middle age to old age, suggesting a steeper deterioration in neural control mechanisms. In contrast, males exhibited a more linear change in motor unit behaviour from young to old age. Sex differences in motor unit firing rates were evident at 30% MVC, with females exhibiting higher firing rates than males in the YG and MA groups (Inglis & Gabriel, 2020). This finding aligns with the hypothesis that females may rely on higher firing rates to compensate for biomechanical differences, such as smaller muscle size and lower force-generating capacity compared to males (Marsala et al., 2025; Olmos et al., 2023; Roos et al., 1997). However, this difference was absent in the OLD group, suggesting that age-related changes in motor unit properties, such as reduced excitability or slower contractile responses, may limit this compensation (Erim et al., 1999; Nelson et al., 1983).

The modulation of firing rates during force ramps was less pronounced in the OLD group, particularly for early-recruited motor units, indicating an age-related decline in the ability to increase firing rates with force demand. Such a pattern was visible in the flattening of the firing rate curves in older adults, particularly in the lower threshold motor unit groups (Figure 4), indicating an age-related limitation in driving early recruited motor units at higher discharge rates. This finding confirms a compromised rate coding mechanism with ageing, where the ability to modulate firing rates according to the recruitment order of motor units is reduced (Heckman & Enoka, 2012), consistently with studies reporting altered discharge patterns in older adults (Erim et al., 1999; Girts et al., 2020; Watanabe et al., 2016), also linked to reductions in both muscle quantity and quality (Hirono et al., 2024). Sex-specific differences further highlight the complexity of this decline. While MA females exhibited greater firing rate modulation than males, this trend reversed in the OLD group, suggesting a steeper decline in females. Finally, the age-related change in modulation is less evident for the late-recruited motor units, perhaps because architectural alterations have a greater impact on the early motor units, which are more actively involved in fine force control (Lecce et al., 2024). Indeed, ageing is also associated with changes in the spatial arrangement of muscle fibres, leading to alterations in muscle architecture, such as reductions in fascicle length and changes in pennation angle (Morse et al., 2005; Narici et al., 2021).

These results indicate sex-specific ageing trajectories, with women maintaining relatively preserved neural function until middle age, followed by a sharper decline—potentially influenced by hormonal changes. Indeed, women face unique challenges such as menopause-related hormonal changes (Al-Azzawi & Palacios, 2009; McNulty et al., 2020) that can influence neuromuscular function (O’Bryan et al., 2025; Sipilä et al., 2015), as fluctuations in estrogenic appear to modulate motor unit behaviour (Jenz et al., 2023; J. Piasecki et al., 2023). In this study, all middle-aged female participants were postmenopausal, with an average age at menopause of 51 ± 4 years and an average of 7 ± 5 years since menopause at the time of enrolment.

Motor unit loss cannot be excluded as potential factor, however literature reports this loss to not be sex-dependent (Jones et al., 2022), leaving other factors as main contributors to the convergence between old males and females.

Another interpretive lens is the fibre-type composition. Males typically have a higher proportion of type II fibres, which are more susceptible to age-related denervation. Evidence suggests a fibre type-specific atrophy (Deschenes, 2004), with type II (fast-twitch) fibres showing a 10–40% size reduction from young to old individuals, while type I (slow-twitch) fibres are relatively preserved (Evans & Lexell, 1995). On one hand this explains the general observed morphological decline, but on the other hand this may explain why men begin to show neural decline earlier, as these fibres undergo remodelling sooner. Conversely, women—who may have a higher prevalence of type I fibres—maintain motor unit firing modulation until middle age, showing a behaviour more similar to the young cohort. The convergence of motor unit behaviour in old age could reflect a shared endpoint of fibre-type homogenization, where denervation outpaces reinnervation, leading to similar fibre composition and neural control strategies in both sexes. The segmentation of motor units by recruitment threshold and the analysis of firing rate modulation provided a nuanced view of these changes. While this approach revealed subtle sex- and age-related differences of low/high threshold motor units, further investigations would require muscle biopsies—which were beyond of the scope of the present work—to confirm this interpretation.

In terms of mechanical output, females demonstrated higher torque variability (CoV) than males at 30% MVC contractions across all age groups, confirming earlier findings in the vastus lateralis and tibialis anterior (Guo et al., 2022, 2023; J. Piasecki et al., 2021). This poorer torque steadiness in females may be attributed to differences in musculotendinous properties (e.g. reduced tendon elasticity (Kubo et al., 2003)) as well as from lower absolute strength (Inglis & Gabriel, 2021). Torque variability also increased with age, particularly in females, indicating an age-related decline in the ability to produce steady force (Castronovo et al., 2018; Enoka & Farina, 2021; Galganski et al., 1993). These effects were more evident at 30% MVC than at 50%, supporting the notion that age-related declines in neuromuscular control are amplified at lower contraction intensities (Castronovo et al., 2018; Oomen & Van Dieën, 2017; J. Piasecki et al., 2021). Females also exhibited greater variability in motor unit firing rates (CoV of firing rates) across all age groups. This may explain the female poorer torque steadiness, and could reflect greater synaptic noise or less effective common input modulation (Orssatto et al., 2022). However, despite poorer torque steadiness, the OLD group showed reduced firing rates variability, particularly at 50% MVC. This may reflect reduced flexibility in motor output and lower discharge modulation capacity in older adults, thus a narrower range would make any small variability disproportionately influential on force output given reduced muscle strength (Enoka & Farina, 2021).

Interestingly, while neural properties converged between sexes in older age, muscle morphology maintained consistent sex differences across all age groups. Males exhibited significantly larger CSA and greater muscle thickness compared to females in all age groups, consistent with previous findings that males generally have greater muscle mass due to androgenic and architectural factors (Guo et al., 2023; Marsala et al., 2025). Our results showed a similar reduction in muscle thickness from YG to OLD (∼4 mm in males and ∼5 mm for females) and also for CSA from MA to OLD (∼2 cm^2^ in both sexes), in contrast with previous evidence that age-related muscle loss may be more pronounced in men (Castronovo et al., 2018; Mitchell et al., 2012; Morse et al., 2005; Watanabe et al., 2016). This suggests that morphological ageing follows a more uniform and sex-preserved trajectory, unlike the neural profile which appears to decouple from morphology in late life.

Finally, physical activity emerged as a critical modulating factor and the results are in line with age-matched data in literature (Wanigatunga et al., 2023). Daily physical activity was similar between YG and MA but reduced in OLD. Interestingly, sex differences were found only in MA and OLD participants where females showed a more active behaviour compared to males, partially in accordance with Belcher et al., 2021 where adult females appeared to have higher activity than males. Although both sexes showed reduced activity levels with age, older females might experience a relatively greater decline (16% vs 10% reduction of active time), which may partly explain their steeper neural deterioration. This supports the hypothesis that neural ageing is more sensitive to behavioural factors, such as activity levels (Jones et al., 2021), than muscle morphology, which may be more genetically or hormonally anchored (Roth, 2012).

## Conclusions

This study provides new insights into the sex- and age-related differences in neuromuscular function by simultaneously assessing muscle morphology, motor unit behaviour, and physical activity across adulthood. The results highlight that while males and females adopt distinct neuromuscular strategies during early and middle adulthood, advanced age leads to a convergence in motor unit behaviour, likely driven by cumulative motor neuron loss, fibre-type homogenization, and impaired rate coding mechanisms. The greater torque variability and higher motor unit firing rates observed in females may reflect compensatory neural strategies to offset biomechanical disadvantages, such as smaller muscle size and lower force-generating capacity. On the other hand, the diminished modulation of firing rates and reduced discharge rates in older adults underscore the progressive decline in motor unit control with ageing. Taken together, these results point to a multifactorial deterioration in motor unit control with ageing, influenced by neural, muscular, hormonal, and biomechanical factors. These findings emphasize the importance of considering sex in the study of neuromuscular ageing and suggest that preserving motor unit recruitment and discharge properties may be critical targets for interventions. Future studies should better understand the mechanisms driving these age- and sex-related differences in neuromuscular function particularly exploring the interplay between muscle morphology, endocrine changes, motor unit behaviour, and the role of exercise training in mitigating their decline, with the goal of developing sex-specific strategies to preserve neuromuscular function in ageing populations. These data will also help to improve physical activity guidelines and pose the basis for more tailored approaches. Finally, longitudinal assessment of these factors along two or more years are warranted to better understand their actual trajectories across life span.

## AUTHOR CONTRIBUTIONS

F.L., R.R, S.P., M.V.F., and A.B. conceptualized the project. M.C. and A.B. conceived the study and designed the experiments. M.C., M.B., M.V.F., A.P. and C.M.B. performed the experiments and collected the data. M.C., M.B., A.P. and C.M.B. analysed the data. M.C., M.B., C.M.B., M.V.F., and A.B interpreted the data results. M.C., and A.B. wrote the original draft. All authors reviewed, contributed, approved the final manuscript, and are accountable for all aspects of the work. All persons designated as authors qualify for authorship, and all those who qualify for authorship are listed.

## ACKNOWLEDGMENTS

The Authors acknowledge the contribution of Federpensionati Coldiretti association in publicizing and promoting the TRAJECTOR-AGE project among its members on a fully free and unconditional basis.

## REFERENCES

Belcher, B. R., Wolff-Hughes, D. L., Dooley, E. E., Staudenmayer, J., Berrigan, D., Eberhardt, M. S., & Troiano, R. P. (2021). US Population-referenced Percentiles for Wrist-Worn Accelerometer-derived Activity. Medicine and Science in Sports and Exercise, 53(11), 2455–2464. 10.1249/MSS.0000000000002726

Brown, W. F. (1972). A method for estimating the number of motor units in thenar muscles and the changes in motor unit count with ageing. Journal of Neurology, Neurosurgery & Psychiatry, 35(6), 845–852. 10.1136/jnnp.35.6.845

Brown, W. F., Strong, M. J., & Snow, R. (1988). Methods for estimating numbers of motor units in biceps-brachialis muscles and losses of motor units with aging. Muscle & Nerve, 11(5), 423–432. 10.1002/mus.880110503

Castronovo, A. M., Mrachacz-Kersting, N., Stevenson, A. J. T., Holobar, A., Enoka, R. M., & Farina, D. (2018). Decrease in force steadiness with aging is associated with increased power of the common but not independent input to motor neurons. Journal of Neurophysiology, 120(4), 1616–1624. 10.1152/jn.00093.2018

Cerone, G. L., Botter, A., & Gazzoni, M. (2019). A Modular, Smart, and Wearable System for High Density sEMG Detection. IEEE Transactions on Biomedical Engineering. 10.1109/tbme.2019.2904398

Cerone, G. L., Giangrande, A., Ghislieri, M., Gazzoni, M., Piitulainen, H., & Botter, A. (2022). Design and validation of a wireless Body Sensor Network for integrated EEG and HD-sEMG acquisitions. IEEE Transactions on Neural Systems and Rehabilitation Engineering. 10.1109/TNSRE.2022.3140220

Del Vecchio, A., Holobar, A., Falla, D., Felici, F., Enoka, R. M., & Farina, D. (2020). Tutorial: Analysis of motor unit discharge characteristics from high-density surface EMG signals. Journal of Electromyography and Kinesiology, 53, 102426. 10.1016/j.jelekin.2020.102426

Delmonico, M. J., Harris, T. B., Visser, M., Park, S. W., Conroy, M. B., Velasquez-Mieyer, P., Boudreau, R., Manini, T. M., Nevitt, M., Newman, A. B., & Goodpaster, B. H. (2009). Longitudinal study of muscle strength, quality, and adipose tissue infiltration12. The American Journal of Clinical Nutrition, 90(6), 1579–1585. 10.3945/ajcn.2009.28047

Deschenes, M. R. (2004). Effects of Aging on Muscle Fibre Type and Size. Sports Medicine, 34(12), 809–824. 10.2165/00007256-200434120-00002

Drey, M., Krieger, B., Sieber, C. C., Bauer, J. M., Hettwer, S., Bertsch, T., Dahinden, P., Mäder, A., Vrijbloed, J. W., Schuster, G., Zollinger, S., Beeler, C., & Unterer, T. (2014). Motoneuron Loss Is Associated With Sarcopenia. Journal of the American Medical Directors Association, 15(6), 435–439. 10.1016/j.jamda.2014.02.002

Enoka, R. M., & Farina, D. (2021). Force Steadiness: From Motor Units to Voluntary Actions. Physiology, 36(2), 114–130. 10.1152/physiol.00027.2020

Erim, Z., Beg, M. F., Burke, D. T., & De Luca, C. J. (1999). Effects of Aging on Motor-Unit Control Properties. Journal of Neurophysiology, 82(5), 2081–2091. 10.1152/jn.1999.82.5.2081

Franchi, M. V., Monti, E., Carter, A., Quinlan, J. I., Herrod, P. J. J., Reeves, N. D., & Narici, M. V. (2019). Bouncing Back! Counteracting Muscle Aging With Plyometric Muscle Loading. Frontiers in Physiology, 10. 10.3389/fphys.2019.00178

Galganski, M. E., Fuglevand, A. J., & Enoka, R. M. (1993). Reduced control of motor output in a human hand muscle of elderly subjects during submaximal contractions. Journal of Neurophysiology, 69(6), 2108–2115. 10.1152/jn.1993.69.6.2108

Girts, R. M., Mota, J. A., Harmon, K. K., MacLennan, R. J., & Stock, M. S. (2020). Vastus lateralis motor unit recruitment thresholds are compressed towards lower forces in older men. Journal of Frailty & Aging, 1–6. 10.14283/jfa.2020.19

Guo, Y., Jones, E. J., Škarabot, J., Inns, T. B., Phillips, B. E., Atherton, P. J., & Piasecki, M. (2024). Common synaptic inputs and persistent inward currents of vastus lateralis motor units are reduced in older male adults. GeroScience, 46(3), 3249–3261. 10.1007/s11357-024-01063-w

Guo, Y., Jones, E. J., Smart, T. F., Altheyab, A., Gamage, N., Stashuk, D. W., Piasecki, J., Phillips, B. E., Atherton, P. J., & Piasecki, M. (2023). Sex disparities of human neuromuscular decline in older humans. 10.1101/2023.06.13.544761

Heckman, C. j., & Enoka, R. M. (2012). Motor Unit. Comprehensive Physiology, 2(4), 2629–2682. 10.1002/j.2040-4603.2012.tb00465.x

Hepple, R. T., & Rice, C. L. (2016). Innervation and neuromuscular control in ageing skeletal muscle. The Journal of Physiology, 594(8), 1965–1978. 10.1113/JP270561

Hirono, T., Takeda, R., Nishikawa, T., Okudaira, M., Kunugi, S., Yoshiko, A., Ueda, S., Yoshimura, A., & Watanabe, K. (2024). Motor unit firing patterns in older adults with low skeletal muscle mass. Archives of Gerontology and Geriatrics, 116, 105151. 10.1016/j.archger.2023.105151

Horwath, O., Moberg, M., Edman, S., Philp, A., & Apró, W. (2025). Ageing leads to selective type II myofibre deterioration and denervation independent of reinnervative capacity in human skeletal muscle. Experimental Physiology, 110(2), 277–292. 10.1113/EP092222

Hunter, S. K., Pereira, H. M., & Keenan, K. G. (2016). The Aging Neuromuscular System and Motor Performance. American Journal of Physiology-Heart and Circulatory Physiology. 10.1152/japplphysiol.00475.2016

Inglis, J. G., & Cabral, H. V. (2025). Unravelling the complexities of neuromuscular function in females throughout the adult lifespan. The Journal of Physiology, n/a(n/a). 10.1113/JP289013

Inglis, J. G., & Gabriel, D. A. (2020). Sex differences in motor unit discharge rates at maximal and submaximal levels of force output. Applied Physiology, Nutrition, and Metabolism, 45(11), 1197–1207. 10.1139/apnm-2019-0958

Inglis, J. G., & Gabriel, D. A. (2021). Sex differences in the modulation of the motor unit discharge rate leads to reduced force steadiness. Applied Physiology, Nutrition, and Metabolism, 46(9), 1065–1072. 10.1139/apnm-2020-0953

Janssen, I., Heymsfield, S. B., & Ross, R. (2002). Low Relative Skeletal Muscle Mass (Sarcopenia) in Older Persons Is Associated with Functional Impairment and Physical Disability. Journal of the American Geriatrics Society, 50(5), 889–896. 10.1046/j.1532-5415.2002.50216.x

Jenz, S. T., Beauchamp, J. A., Gomes, M. M., Negro, F., Heckman, C. J., & Pearcey, G. E. P. (2023). Estimates of persistent inward currents in lower limb motoneurons are larger in females than in males. Journal of Neurophysiology, 129(6), 1322–1333. 10.1152/jn.00043.2023

Jones, E. J., Chiou, S.-Y., Atherton, P. J., Phillips, B. E., & Piasecki, M. (2022). Ageing and exercise-induced motor unit remodelling. The Journal of Physiology, 600(8), 1839–1849. 10.1113/JP281726

Jones, E. J., Piasecki, J., Ireland, A., Stashuk, D. W., Atherton, P. J., Phillips, B. E., McPhee, J. S., & Piasecki, M. (2021). Lifelong exercise is associated with more homogeneous motor unit potential features across deep and superficial areas of vastus lateralis. GeroScience, 43(4), 1555–1565. 10.1007/s11357-021-00356-8

Karas, M., Schrack, J., & Urbanek, J. (2021). R: arctools: Processing and physical activity summaries of… [Computer software]. https://search.r-project.org/CRAN/refmans/arctools/html/arctools-package.html

Klass, M., Baudry, S., & Duchateau, J. (2008). Age-related decline in rate of torque development is accompanied by lower maximal motor unit discharge frequency during fast contractions. Journal of Applied Physiology (Bethesda, Md.: 1985), 104(3), 739–746. 10.1152/japplphysiol.00550.2007

Koster, A., Shiroma, E. J., Caserotti, P., Matthews, C. E., Chen, K. Y., Glynn, N. W., & Harris, T. B. (2016). Comparison of Sedentary Estimates between activPAL and Hip- and Wrist-Worn ActiGraph. Medicine and Science in Sports and Exercise, 48(8), 1514–1522. 10.1249/MSS.0000000000000924

Larsson, L., Degens, H., Li, M., Salviati, L., Lee, Y. il, Thompson, W., Kirkland, J. L., & Sandri, M. (2019). Sarcopenia: Aging-Related Loss of Muscle Mass and Function. Physiological Reviews, 99(1), 427–511. 10.1152/physrev.00061.2017

Lauretani, F., Maggio, M., Pilotto, A. M., Ansaldo, M., Brusco, C. M., Carbonaro, M., Amendola, C., Nabacino, M., Testa, C., Ciuni, A., Sverzellati, N., Zucchini, I., Salvi, M., Mastropietro, A., Re, R., Botter, A., Franchi, M. V., & Porcelli, S. (2025). The Trajectories of Neuromuscular Aging (TRAJECTOR-AGE Clinical Trial): Study Rationale and Methodological Protocol. Journal of the American Geriatrics Society, n/a(n/a). 10.1111/jgs.70005

Lecce, E., Conti, A., Nuccio, S., Felici, F., & Bazzucchi, I. (2024). Characterising sex-related differences in lower- and higher-threshold motor unit behaviour through high-density surface electromyography. Experimental Physiology, 109(8), 1317–1329. 10.1113/EP091823

Lexell, J., & Downham, D. Y. (1991). The occurrence of fibre-type grouping in healthy human muscle: A quantitative study of cross-sections of whole vastus lateralis from men between 15 and 83 years. Acta Neuropathologica, 81(4), 377–381. 10.1007/BF00293457

Lexell, J., Taylor, C. C., & Sjöström, M. (1988). What is the cause of the ageing atrophy?: Total number, size and proportion of different fiber types studied in whole vastus lateralis muscle from 15- to 83-year-old men. Journal of the Neurological Sciences, 84(2), 275–294. 10.1016/0022-510X(88)90132-3

Long, J. S., & Pavalko, E. K. (2004). The Life Course of Activity Limitations: Exploring Indicators of Functional Limitations Over Time. Journal of Aging and Health, 16(4), 490–516. 10.1177/0898264304265776

Lulic-Kuryllo, T., & Inglis, J. G. (2022). Sex differences in motor unit behaviour: A review. Journal of Electromyography and Kinesiology, 66(July), 102689. 10.1016/j.jelekin.2022.102689

Mani, D., M, A. A., D, H. L., M, V. T., Botter, A., & M, E. R. (2018). Motor unit activity, force steadiness, and perceived fatigability are correlated with mobility in older adults. JOURNAL OF NEUROPHYSIOLOGY, 120, 1988–1997. 10.1152/jn.00192.2018

Marsala, M. J., Kells, A. M., & Christie, A. D. (2025). Sex-related differences in motor unit firing rate and pennation angle. Applied Physiology, Nutrition, and Metabolism, 50, 1–9. 10.1139/apnm-2024-0202

Merletti, R., & Muceli, S. (2019). Tutorial. Surface EMG detection in space and time: Best practices. Journal of Electromyography and Kinesiology, 49(August), 102363. 10.1016/j.jelekin.2019.102363

Mitchell, W. K., Atherton, P. J., Williams, J., Larvin, M., Lund, J. N., & Narici, M. (2012). Sarcopenia, Dynapenia, and the Impact of Advancing Age on Human Skeletal Muscle Size and Strength; a Quantitative Review. Frontiers in Physiology, 3. 10.3389/fphys.2012.00260

Monti, E., Franchi, M. V., Badiali, F., Quinlan, J. I., Longo, S., & Narici, M. V. (2020). The Time-Course of Changes in Muscle Mass, Architecture and Power During 6 Weeks of Plyometric Training. Frontiers in Physiology, 11, 946. 10.3389/fphys.2020.00946

Morse, C. I., Thom, J. M., Reeves, N. D., Birch, K. M., & Narici, M. V. (2005). In vivo physiological cross-sectional area and specific force are reduced in the gastrocnemius of elderly men. Journal of Applied Physiology (Bethesda, Md.: 1985), 99(3), 1050–1055. 10.1152/japplphysiol.01186.2004

Narici, M., McPhee, J., Conte, M., Franchi, M. V., Mitchell, K., Tagliaferri, S., Monti, E., Marcolin, G., Atherton, P. J., Smith, K., Phillips, B., Lund, J., Franceschi, C., Maggio, M., & Butler-Browne, G. S. (2021). Age-related alterations in muscle architecture are a signature of sarcopenia: The ultrasound sarcopenia index. Journal of Cachexia, Sarcopenia and Muscle, 12(4), 973–982. 10.1002/jcsm.12720

Nelson, R. M., Soderberg, G. L., & Urbscheit, N. L. (1983). Comparison of skeletal muscle motor unit discharge characteristics in young and aged humans. Archives of Gerontology and Geriatrics, 2(3), 255–264. 10.1016/0167-4943(83)90029-8

Nelson, R. M., Soderberg, G. L., & Urbscheit, N. L. (1984). Alteration of Motor-Unit Discharge Characteristics in Aged Humans. Physical Therapy, 64(1), 29–34. 10.1093/ptj/64.1.29

Nishikawa, Y., Watanabe, K., Holobar, A., Kitamura, R., Maeda, N., & Hyngstrom, A. S. (2024). Sex differences in laterality of motor unit firing behavior of the first dorsal interosseous muscle in strength-matched healthy young males and females. European Journal of Applied Physiology, 124(7), 1979–1990. 10.1007/s00421-024-05420-7

O’Bryan, S. J., Critchlow, A., Fuchs, C. J., Hiam, D., & Lamon, S. (2025). The contribution of age and sex hormones to female neuromuscular function across the adult lifespan. The Journal of Physiology, n/a(n/a). 10.1113/JP287496

Olmos, A. A., Sterczala, A. J., Parra, M. E., Dimmick, H. L., Miller, J. D., Deckert, J. A., Sontag, S. A., Gallagher, P. M., Fry, A. C., Herda, T. J., & Trevino, M. A. (2023). Sex-related differences in motor unit behavior are influenced by myosin heavy chain during high- but not moderate-intensity contractions. Acta Physiologica, 239(1), e14024. 10.1111/apha.14024

Orssatto, L. B. R., Blazevich, A. J., & Trajano, G. S. (2023). Ageing reduces persistent inward current contribution to motor neurone firing: Potential mechanisms and the role of exercise. The Journal of Physiology, 601(17), 3705–3716. 10.1113/JP284603

Orssatto, L. B. R., Fernandes, G. L., Blazevich, A. J., & Trajano, G. S. (2022). Facilitation–inhibition control of motor neuronal persistent inward currents in young and older adults. The Journal of Physiology, 600(23), 5101–5117. 10.1113/JP283708

Parra, M. E., Sterczala, A. J., Miller, J. D., Trevino, M. A., Dimmick, H. L., & Herda, T. J. (2020). Sex-related differences in motor unit firing rates and action potential amplitudes of the first dorsal interosseous during high-, but not low-intensity contractions. Experimental Brain Research, 238(5), 1133–1144. 10.1007/s00221-020-05759-1

Piasecki, J., Inns, T. B., Bass, J. J., Scott, R., Stashuk, D. W., Phillips, B. E., Atherton, P. J., & Piasecki, M. (2021). Influence of sex on the age-related adaptations of neuromuscular function and motor unit properties in elite masters athletes. The Journal of Physiology, 599(1), 193–205. 10.1113/JP280679

Piasecki, M., Ireland, A., Piasecki, J., Stashuk, D. W., Swiecicka, A., Rutter, M. K., Jones, D. A., & McPhee, J. S. (2018). Failure to expand the motor unit size to compensate for declining motor unit numbers distinguishes sarcopenic from non-sarcopenic older men. The Journal of Physiology, 596(9), 1627–1637. 10.1113/JP275520

Roos, M. R., Rice, C. L., & Vandervoort, A. A. (1997). Age-related changes in motor unit function. Muscle & Nerve, 20(6), 679–690. 10.1002/(SICI)1097-4598(199706)20:6%253C679::AID-MUS4%253E3.0.CO;2-5

Roth, S. M. (2012). Genetic aspects of skeletal muscle strength and mass with relevance to sarcopenia. BoneKEy Reports, 1, 58. 10.1038/bonekey.2012.58

Sarto, F., Franchi, M. V., McPhee, J. S., Stashuk, D. W., Paganini, M., Monti, E., Rossi, M., Sirago, G., Zampieri, S., Motanova, E. S., Valli, G., Moro, T., Paoli, A., Bottinelli, R., Pellegrino, M. A., De Vito, G., Blau, H. M., & Narici, M. V. (2024). Neuromuscular impairment at different stages of human sarcopenia. Journal of Cachexia, Sarcopenia and Muscle, 15(5), 1797–1810. 10.1002/jcsm.13531

Sundberg, C. W. (2025). Physiology of ageing skeletal muscle and the protective effects of exercise. The Journal of Physiology, 603(1), 3–6. 10.1113/JP287926

Tieland, M., Trouwborst, I., & Clark, B. C. (2018). Skeletal muscle performance and ageing. Journal of Cachexia, Sarcopenia and Muscle, 9(1), 3–19. 10.1002/jcsm.12238

Wanigatunga, A. A., Liu, F., Urbanek, J. K., Wang, H., Di, J., Zipunnikov, V., Cai, Y., Dougherty, R. J., Simonsick, E. M., Ferrucci, L., & Schrack, J. A. (2023). Wrist-Worn Accelerometry, Aging, and Gait Speed in the Baltimore Longitudinal Study of Aging. Journal of Aging and Physical Activity, 31(3), 408–416. 10.1123/japa.2022-0156

Watanabe, K., Holobar, A., Kouzaki, M., Ogawa, M., Akima, H., & Moritani, T. (2016). Age-related changes in motor unit firing pattern of vastus lateralis muscle during low-moderate contraction. AGE, 38(3), 48. 10.1007/s11357-016-9915-0

Wilkinson, D. J., Piasecki, M., & Atherton, P. J. (2018). The age-related loss of skeletal muscle mass and function: Measurement and physiology of muscle fibre atrophy and muscle fibre loss in humans. Ageing Research Reviews, 47, 123–132. 10.1016/j.arr.2018.07.005

Yacyshyn, A. F., Mohammadalinejad, G., Afsharipour, B., Duchcherer, J., Bashuk, J., Bennett, D. J., Negro, F., Quinlan, K. A., & Gorassini, M. A. (2025). Sex-related differences in motoneuron firing behavior during typical development. Journal of Neurophysiology, 133(4), 1307–1319. 10.1152/jn.00505.2024

